# Biofilm Formation by *Pseudomonas aeruginosa* in Medium Supplemented with Adult Bovine Serum and the Anti-Biofilm Activity of β-Lactam Antibiotics

**DOI:** 10.1101/2025.09.14.676195

**Authors:** Junichi Mitsuyama, Kyoko Asano, Yoshitomo Morinaga

## Abstract

*Pseudomonas aeruginosa* is a major opportunistic pathogen known for its ability to form biofilms in vivo, which contributes significantly to its resistance to antimicrobial treatment. In this study, we examined the biofilm-forming capacity of a clinical isolate cultured in medium supplemented with 30% adult bovine serum (ABS). Under these host-mimicking conditions, mature biofilms developed on microplate surfaces within 8 hours. Among the β-lactam antibiotics evaluated, piperacillin and cefoperazone exhibited potent anti-biofilm activity at concentrations as low as 1/8,192 to 1/256 of their respective minimum inhibitory concentrations (MICs). In contrast, ceftazidime and meropenem required significantly higher concentrations (approximately 1/16 MIC) to produce similar effects. The anti-biofilm activity of piperacillin and cefoperazone was abolished when ABS was defatted with n-hexane or when magnesium sulfate was added to the medium, suggesting that a lipid-dependent membrane permeabilization mechanism may be involved. These findings highlight the utility of host-mimicking conditions in evaluating the activity of antibiotics against biofilms and suggest that certain β-lactam agents may have previously unrecognized anti-biofilm effects mediated through serum-associated components.

## Introduction

*Pseudomonas aeruginosa* is a prominent member of the ESKAPE group of pathogens—comprising *Enterococcus faecium*, *Staphylococcus aureus*, *Klebsiella pneumoniae*, *Acinetobacter baumannii*, *P. aeruginosa*, and *Enterobacter* species—which are collectively recognized for their high virulence and extensive antimicrobial resistance [1]. This opportunistic Gram-negative bacillus is a leading cause of healthcare-associated infections, including ventilator-associated pneumonia, catheter-related bloodstream infections, gastrointestinal infections, dermatitis, folliculitis, otitis externa, and surgical site infections, particularly in immunocompromised individuals [2].

A key factor contributing to the persistence and therapeutic recalcitrance of *P. aeruginosa* is its ability to form biofilms—organized communities of bacteria embedded within a self-produced extracellular polymeric substance (EPS) matrix [3,4]. Within these biofilms, bacterial cells exhibit phenotypic heterogeneity characterized by reduced metabolic activity and microenvironment-dependent dormancy [5], altered transcriptional profiles with upregulation of stress response and stationary-phase regulons [6], and significantly increased tolerance to antimicrobial agents and host immune mechanisms [6,7]. The biofilm matrix not only acts as a physical barrier that limits antibiotic penetration but also actively sequesters antimicrobial compounds—particularly cationic agents—thereby diminishing their therapeutic efficacy [8].

Although the biology of *P. aeruginosa* biofilms has been extensively studied, most experimental investigations have utilized artificial, modified, or minimal culture media [9,10]. While such systems offer convenience, reproducibility, and cost-effectiveness, they fail to fully recapitulate the complex in vivo microenvironment, including factors such as nutrient gradients, immune-mediated pressures, and serum-derived components. Consequently, there is a critical need for physiologically relevant *in vitro* models that incorporate host-derived elements, such as serum, to more accurately mimic *in vivo* biofilm behavior.

Hammond *et al.* [11] previously demonstrated that both bovine and human serum suppress biofilm formation by the laboratory reference strain *P. aeruginosa* PAO1. However, the extensive genetic and phenotypic diversity observed among clinical isolates suggests that some strains may retain the capacity to form mature biofilms even in serum-rich environments.

In the present study, we identified a clinical isolate of *P. aeruginosa* capable of robust biofilm formation in the presence of adult bovine serum (ABS). Using this strain, we investigated the anti-biofilm activity of four β-lactam antibiotics—piperacillin (PIPC), cefoperazone (CPZ), ceftazidime (CAZ), and meropenem (MEPM)—under ABS-enriched conditions. Notably, PIPC and CPZ, both of which contain a 2,3-diketopiperazine (DKP) structural motif, exhibited pronounced anti-biofilm effects at concentrations markedly lower than their respective minimum inhibitory concentrations (MICs), as determined according to CLSI guidelines. Interestingly, other DKP-containing analogs lacking structural similarity to PIPC and CPZ failed to reproduce this effect, suggesting that the observed activity is not solely attributable to the presence of the DKP moiety, but likely involves a more nuanced interplay of molecular structure and physicochemical properties.

This study aims to elucidate the mechanisms underpinning serum-facilitated biofilm formation by *P. aeruginosa* and to characterize the context-dependent anti-biofilm efficacy of selected β-lactam antibiotics under conditions that more closely reflect the host environment.

## Materials and Methods

### Materials

Piperacillin sodium (PIPC) was obtained from FUJIFILM Toyama Chemical Co.,Ltd (Tokyo, Japan). Cefoperazone (CPZ), ceftazidime (CAZ), and meropenem (MEPM) were purchased from Tokyo Chemical Industry Co., Ltd (Tokyo, Japan). Vancomycin hydrochloride was purchased from FUJIFILM Wako Pure Chemical Corp (Osaka, Japan).

Compounds containing the 2,3-diketopiperazine (DKP) moiety—(R)-2-(4-ethyl-2,3-dioxopiperazine-1-carboxamido)-2-phenylacetic acid (a known impurity of PIPC) and (R)-(-)-α-(((4-ethyl-2,3-dioxo-1-piperazinyl)carbonyl)amino)-4-hydroxybenzenea cetic acid (a known impurity of CPZ)—were also obtained from Tokyo Chemical Industry Co., Ltd. Ethyl 4-ethyl-2,3-dioxopiperazine-1-carboxylate and 1-acetyl-4-ethylpiperazine-2,3-dione were synthesized by NARD Institute, Ltd (Hyogo, Japan).

### Bacterial Strains and Culture Conditions

Five clinical isolates of *P. aeruginosa* were used in this study. The reference strain PAO1 and two isolates (strain S3 and S6) were kindly provided by Nagasaki University, while two additional clinical isolates were obtained from Toyama University Hospital. All bacterial strains were suspended in Dulbecco’s Phosphate-Buffered Saline (DPBS(-); Thermo Fisher Scientific, Oxoid, Basingstoke, UK) supplemented with 50% (w/v) glycerol and stored at −80 °C.

Prior to experimentation, bacterial cultures were initiated by inoculating a loopful of each strain into 5 mL of cation-adjusted Mueller–Hinton broth (CAMHB; BD Difco, Detroit, MI, USA), followed by incubation at 37 °C with shaking at 200 rpm for 24 hours. The resulting cultures had an initial cell density of 7 × 10⁸ to 1 × 10⁹ CFU/mL, confirmed via colony counting on LB agar plates (LBA; Difco, Sparks, MD, USA). For biofilm assays, bacterial suspensions were diluted 1:1000 in adult bovine serum (ABS) medium to achieve a final concentration of 7 × 10^5^ to 1 ×10^6^ CFU/mL. All experiments were performed in triplicate using standardized inocula.

The medium for biofilm formation was prepared as a mixture of 5× salt solution (3 g Na_2_HPO_4_, 1.5 g KH_2_PO_4_, 0.25 g NaCl per 100 mL), adult bovine serum (ABS; Sigma-Aldrich, St. Louis, MO, USA), and sterile water in a ratio of 2:3:5, with minor modifications based on the method described by Hammond *et al*. [11].

### Minimum Inhibitory Concentration (MIC) Determination

MICs of the tested antibiotics against *P. aeruginosa* were determined using the standard broth microdilution method in accordance with Clinical and Laboratory Standards Institute (CLSI) guidelines [12] .

### Quantification of Biofilm Formation by Crystal Violet Assay

Overnight cultures grown in CAMHB at 37 °C were diluted 1:100 in sterile saline. Five microliters of this diluted suspension were added to each well of a polystyrene 96-well microtiter plate containing serial two-fold dilutions of the test antibiotics in biofilm medium. Plates were incubated overnight at 37 °C.

Following incubation, wells were gently washed three times with sterile distilled water to remove non-adherent cells. Adherent biofilms were stained with 0.5% crystal violet for 30 minutes at room temperature. Excess stain was removed by triple washing with distilled water, followed by air drying. The stained biofilm was solubilized with ethanol, and absorbance was measured at 600 nm using a microplate reader (GloMax® Discover GM3000; Promega, Madison, USA). Each condition was evaluated in triplicate. Biofilm formation was quantified by subtracting the absorbance of blank wells, and the 50% biofilm inhibitory concentration (BFIC₅₀) was defined as the lowest drug concentration that reduced OD₆₀₀ by at least 50% compared to untreated controls.

### Determination of Planktonic, Total, and Biofilm-associated Cell Counts

After biofilm formation, planktonic cells were quantified by collecting 10 µL of the culture supernatant, followed by serial dilution in sterile saline and plating on LBA plates. Plates were incubated at 37 °C for 24 hours.

To determine total viable cells (planktonic + biofilm-associated), the wells were vigorously homogenized using sterile small cotton swabs. A 10 µL aliquot of the homogenate was diluted and plated as described above.

For quantification of biofilm-associated cells, wells were washed three times with sterile saline to remove planktonic cells. Residual adherent biofilms were resuspended in 100 µL of saline using sterile small cotton swabs. Ten microliters of the suspension were serially diluted and plated on LBA for CFU enumeration after 24-hour incubation at 37 °C (Fig.1).

**Fig. 1.**
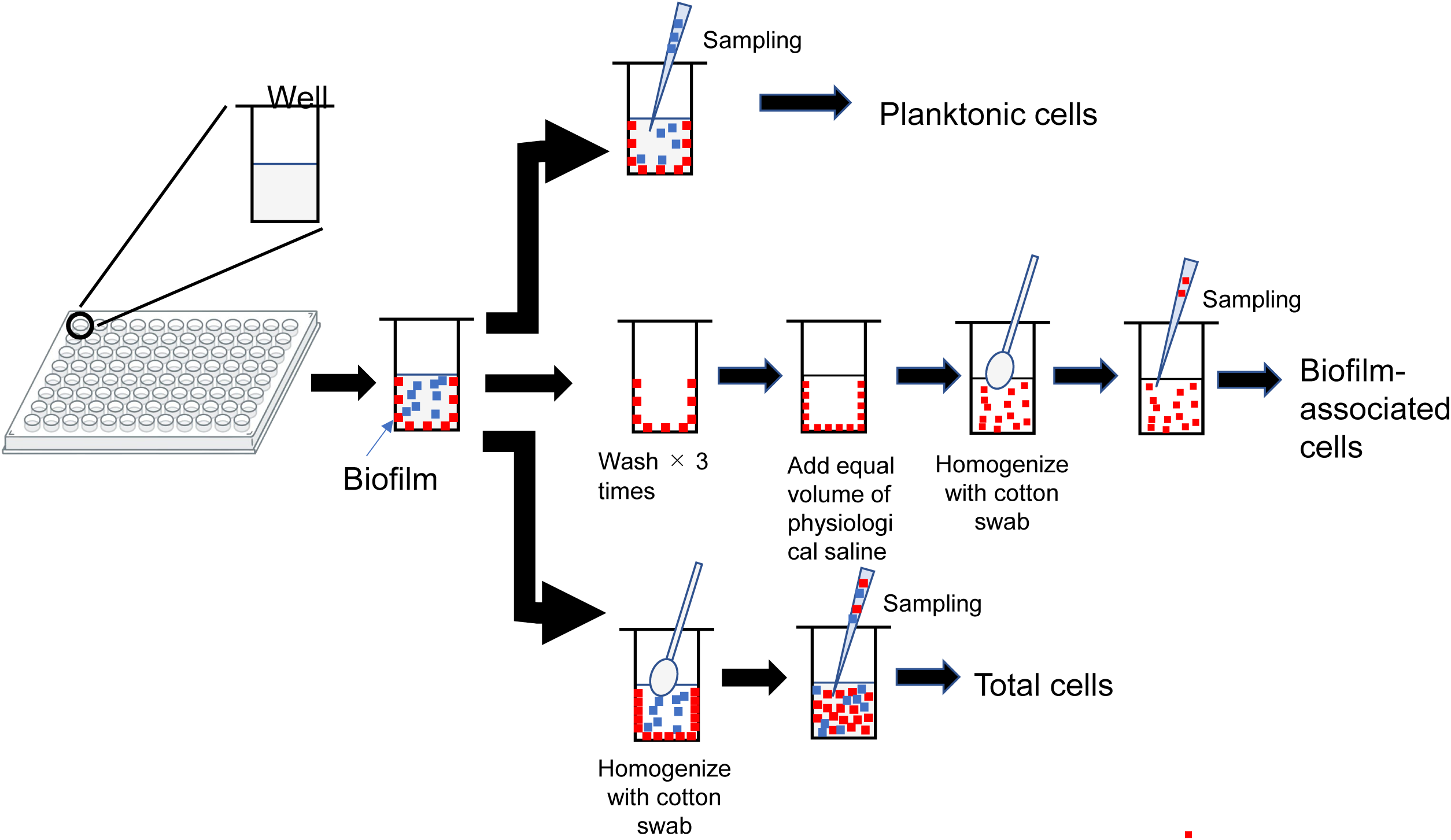
Schematic representation of CFU determination for planktonic, total, and biofilm-associated cells from a 96-well plate.

### Microscopic Observation of Biofilm Formation

An overnight culture of *P. aeruginosa* strain S3 was cultivated in cation-adjusted Mueller–Hinton broth (CAMHB) at 37 °C. The culture was subsequently diluted with ABS medium to achieve a final bacterial concentration of 1 × 10^6^ CFU/mL and cultured in glass-bottomed dish (Matsunami Glass Industry Co., Ltd., Osaka, Japan) at 37°C under static conditions for 8 hours. Prior to imaging by fluorescence microscopy, biofilms were stained using the Live/Dead™ BacLight™ Bacterial Viability Kit (Invitrogen™, Thermo Fisher Scientific, Waltham, MA, USA). Two-dimensional and three-dimensional biofilm structures were subsequently visualized using fluorescence microscopy (BZ-X800®, KEYENCE Corporation, Osaka, Japan).

### Statistical analysis

The data were analyzed using the GraphPad Prism (version 8.4.3) software program (GraphPad Software, La Jolla, CA, USA) and expressed as the mean ± standard deviation (S.D.). The statistical significance of the data was determined by an analysis of variance (ANOVA) test followed by the Dunnet Test.

## Results

### Biofilm Formation in Adult Bovine Serum (ABS)

Among several clinical isolates of *P. aeruginosa*, strains 263, S3, S6, and P-174 exhibited robust biofilm formation in medium supplemented with 30% adult bovine serum (ABS). In contrast, the reference strain PAO1 displayed only minimal biofilm formation under identical conditions (Fig. 2).

**Fig. 2.**
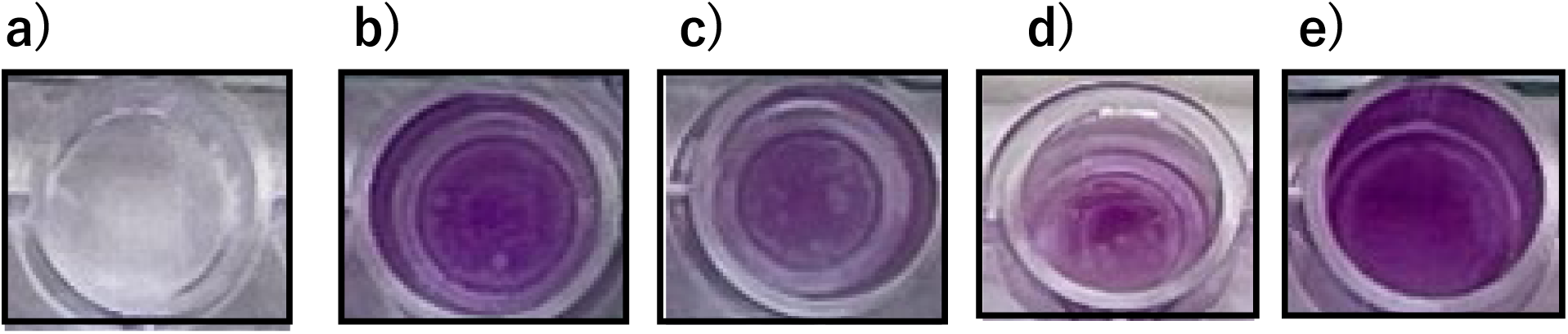
Biofilm formation of *P. aeruginosa* PAO1 and clinical isolates in ABS medium. a) PAO1, b) S3, c) S6, d) 263, e) P-174. Biofilms were developed in 96-well flat-bottom microplates incubated at 37°C for 24 hours in ABS medium. After incubation, wells were washed with sterile distilled water and stained with 0.5% crystal violet for 30 minutes at room temperature. Excess stain was removed by washing, and plates were air-dried.

Fig.3 panel a) illustrates the number of planktonic (supernatant) and biofilm-associated *P. aeruginosa* strain S3 cells cultured in ABS medium. During the first 4 hours of incubation, the number of biofilm-associated cells remained less than 1% of the viable planktonic cell population. However, by 8 hours, the number of biofilm-associated cells surpassed that of the planktonic cells. After 24 hours of incubation, the viable cell counts in the planktonic and biofilm-associated populations were approximately equivalent.

**Fig. 3.**
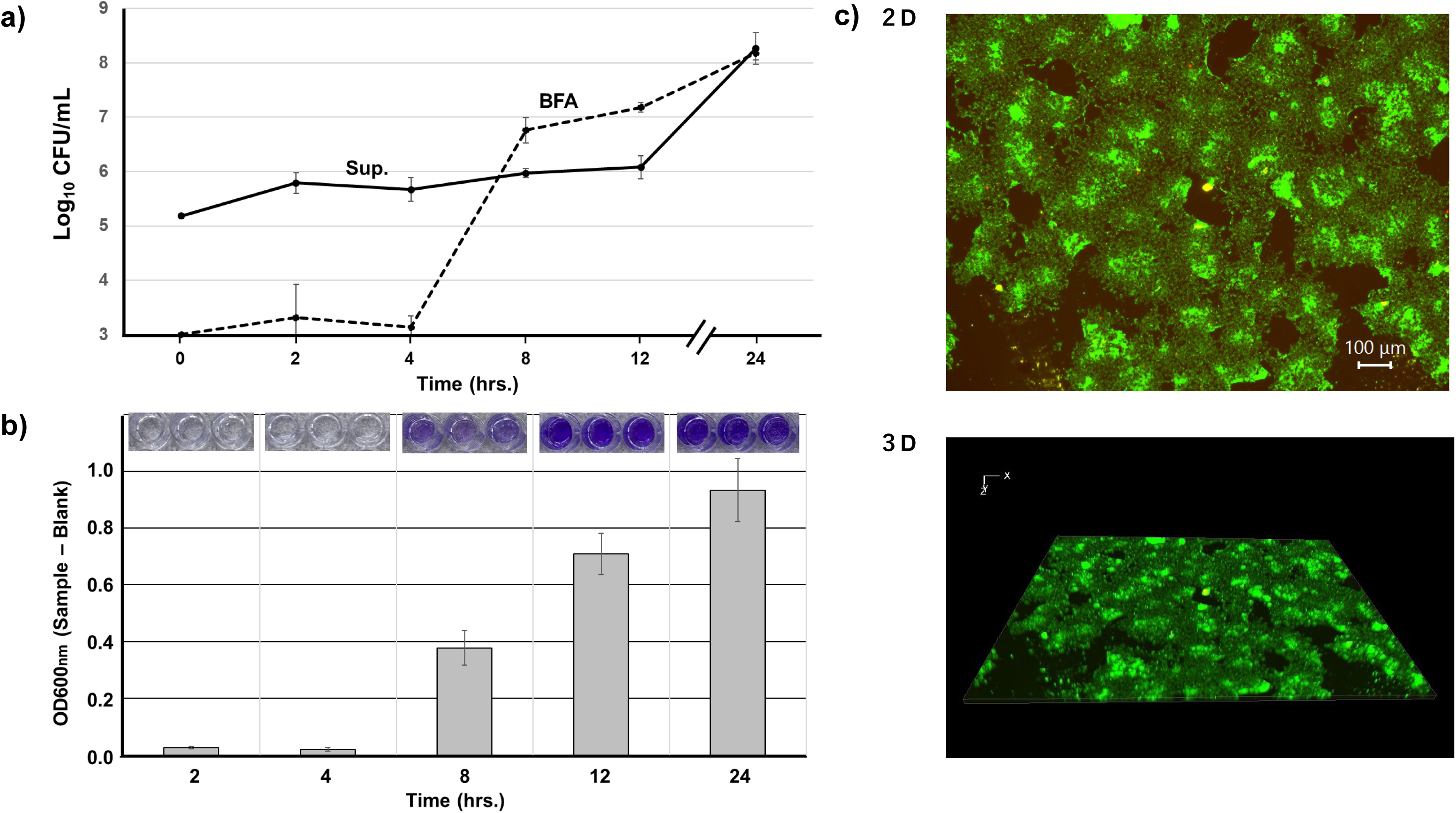
Biofilm formation process of *P. aeruginosa* strain S3. a) Number of cells in supernatant and biofilm-associated cells in ABS medium. Sup.: planktonic cells in culture broth, BFA: biofilm-associated cells. Solid and dot line show the CFU/mL of planktonic and biofilm-associated cells, respectively. Each experiment was evaluated in triplicate, and expressed as average ±S.D. b) Crystal violet staining at the indicated time points and absorbance of OD_600_nm in each time point. c) Imaging for biofilm formation in ABS medium in a glass-bottomed dish using fluorescence microscope. 2D and 3D images at 8 hours.

Biofilm formation, as evaluated by crystal violet staining, is shown in Fig.3 panel b), with the corresponding optical density measurements at 600 nm (OD₆₀₀). Consistent with the trend observed in viable cell counts, minimal biofilm accumulation was detected during the initial 4 hours of incubation. By 8 hours, however, discernible staining patterns characteristic of biofilm formation emerged, followed by a progressive, time-dependent increase in staining intensity. These findings suggest that biofilm development is closely associated with a gradual increase in the number of viable cells embedded within the biofilm matrix. Fig. 3 panel C depicts both two-dimensional and three-dimensional images of the biofilm architecture after 8 hours of incubation. Biofilms did not form as a homogeneous layer but developed as irregularly distributed mushroom-shaped microcolonies on the solid surface. Over the course of the culture period, these microcolonies increased in both frequency and size, eventually fusing to form larger, more complex biofilm structures.

### Anti-Biofilm Activity of β-Lactam Antibiotics

The minimum inhibitory concentrations (MICs) of PIPC, CPZ, CAZ, and MEPM against the test strains are summarized in Table 1. Strains S3 and S6 were resistant to PIPC and CPZ but remained susceptible to CAZ and MEPM, whereas strains 263 and P-174, identified as carbapenemase producers, demonstrated resistance to all agents tested.

**Table 1.**
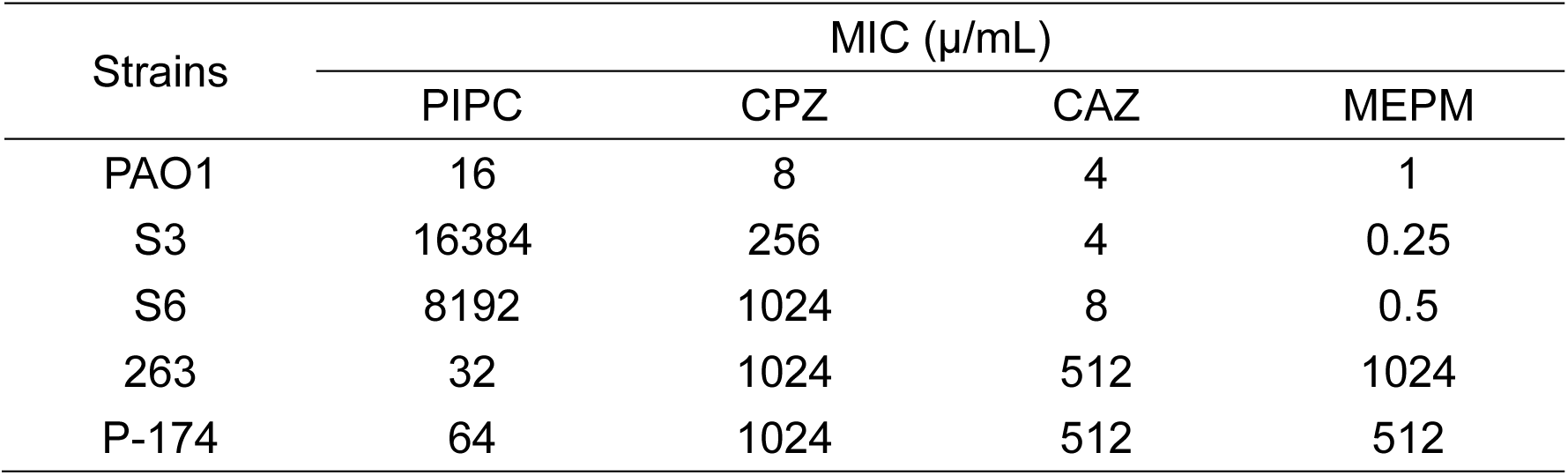
MICs of PIPC, CPZ, CAZ, and MEPM against clinical *P. aeruginosa* isolates.

| Strains | MIC ($\mu$/mL) | | | |
| --- | --- | --- | --- | --- |
|  | PIPC | CPZ | CAZ | MEPM |
| PAO1 | 16 | 8 | 4 | 1 |
| S3 | 16384 | 256 | 4 | 0.25 |
| S6 | 8192 | 1024 | 8 | 0.5 |
| 263 | 32 | 1024 | 512 | 1024 |
| P-174 | 64 | 1024 | 512 | 512 |

**Table 2.** MIC and BFIC₅₀ values of 2,3-DKP analogs. All compounds exhibited MICs and BFIC₅₀ values >1024 µg/mL.

| Compound | MIC (µg/mL) | BFIC<sub>50</sub> (µg/mL) |
| --- | --- | --- |
| PIPC | 16384 | 2 |
| CPZ | 256 | 1 |
| C | >1024 | >1024 |
| D | >1024 | >1024 |
| E | >1024 | >1024 |
| F | >1024 | >1024 |

\*: Biofilm inhibitory concentration 50%

Biofilm inhibition was evaluated using strain S3, which forms stable biofilms in ABS medium. CAZ and MEPM suppressed biofilm formation at concentrations as low as 1/16 of their MICs. Notably, PIPC and CPZ inhibited biofilm formation at concentrations between 1/8192 and 1/256 of their MICs. The biofilm formation inhibitory concentration 50% (BFIC₅₀) values were: PIPC 2 µg/mL, CPZ 1 µg/mL, CAZ 0.25 µg/mL, and MEPM 0.063 µg/mL (Fig.4).

**Fig. 4.**
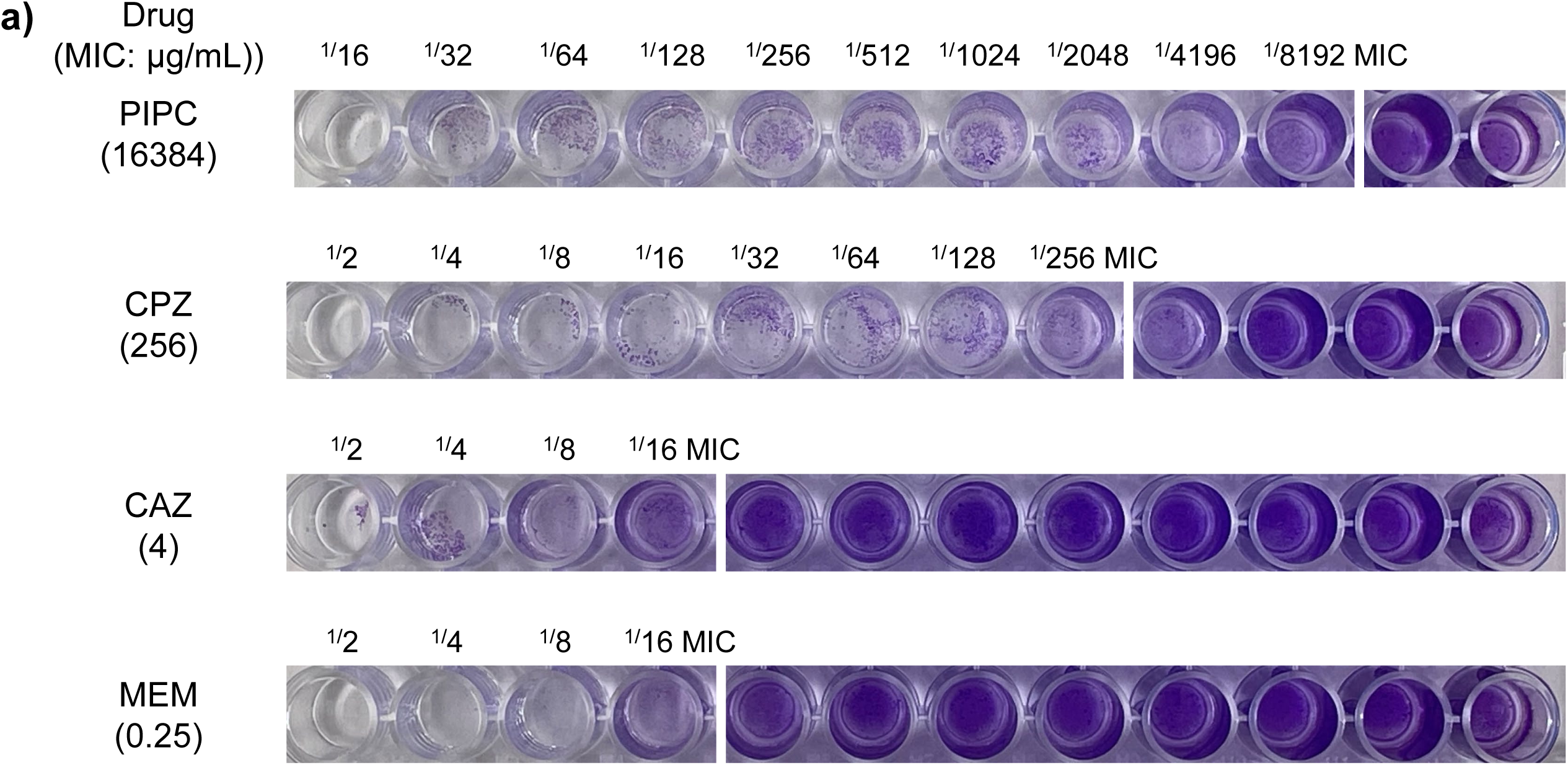
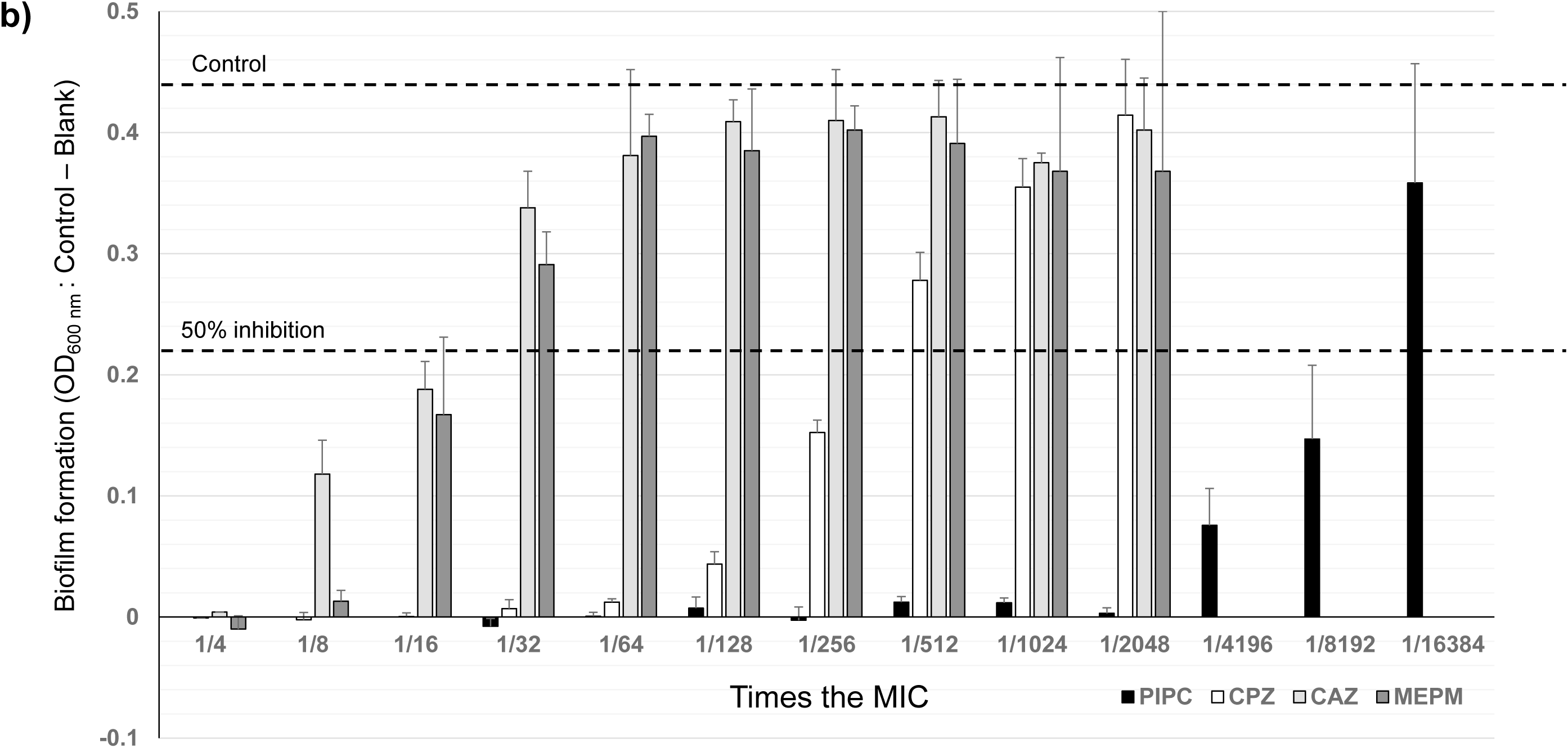
a) Dose-dependent inhibition of biofilm formation by sub-MIC concentrations of each antibiotic in *P. aeruginosa* strain S3. White line indicates the BFIC_50_ for each agent. BFIC₅₀ was defined as the lowest concentration yielding a ≥50% reduction in OD_600_ compared to control. b) Quantification via absorbance measurement (OD_600_) post crystal violet staining and ethanol solubilization. Each experiment was evaluated in triplicate, and expressed as average ±S.D.

## 3. Evaluation of 2,3-Diketopiperazine Derivatives

Given the shared 2,3-diketopiperazine (2,3-DKP) moiety in PIPC and CPZ, we hypothesized that this structure might contribute to biofilm inhibition. Compounds C and D (impurities of PIPC and CPZ) and Compounds E and F (comprising the terminal 2,3-DKP structure) were evaluated (Fig. 5).

**Fig. 5.**
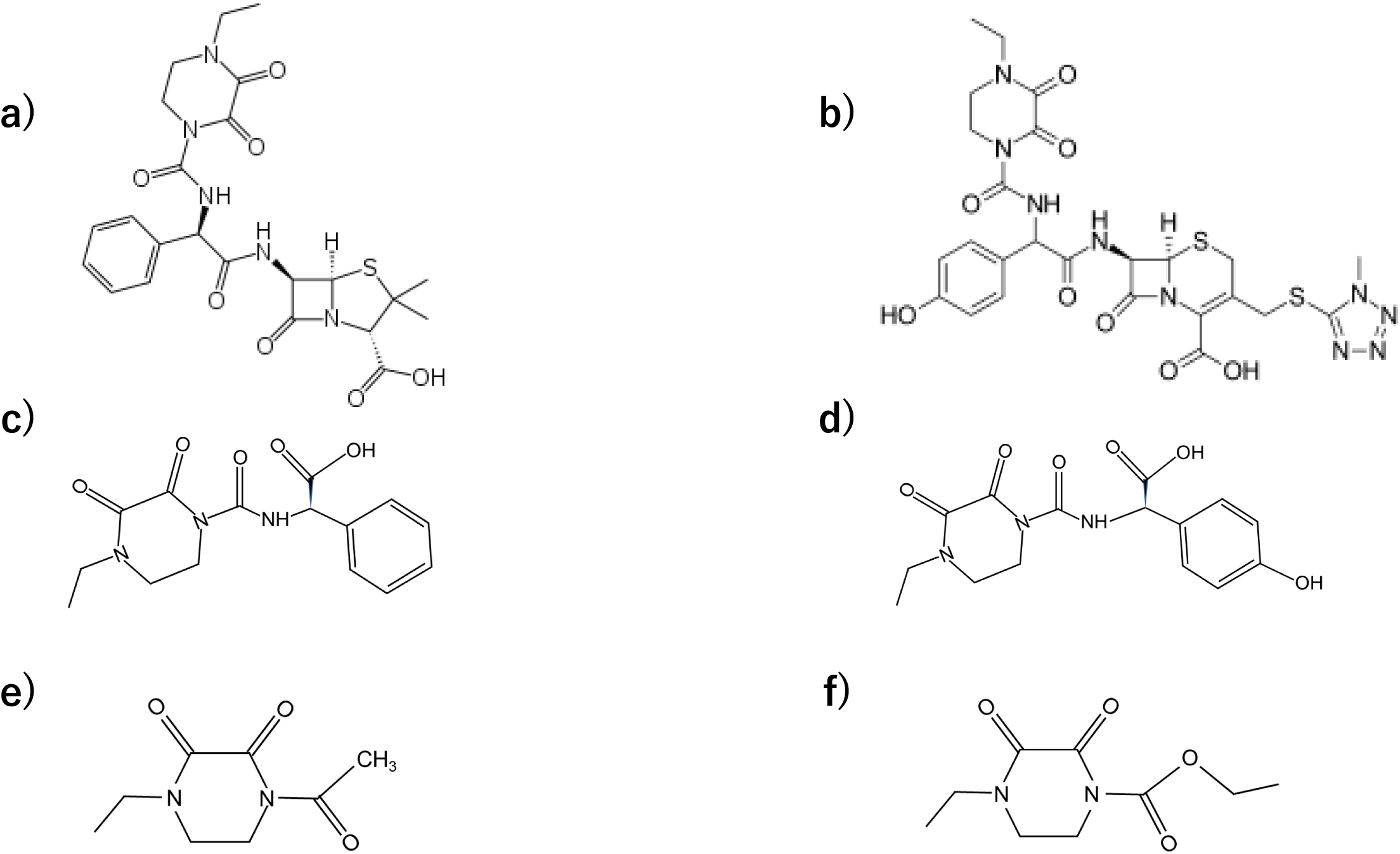
Chemical structure of 2,3-diketopiperazine derivatives a): piperacillin, b): cefoperazone, c) compound C: (2*R*)-2-[(4-ethyl-2,3-dioxopiperazine-1-carbonyl)amino]-2-phenylacetic acid, d) compound D: (2*R*)-2-[(4-ethyl-2,3-dioxopiperazine-1-carbonyl)amino]-2-(4-hydroxyphenyl) acetic acid, e) compound E: 1-acetyl-4-ethylpiperazine-2,3-dione, f) compound F: ethyl 4-ethyl-2,3-dioxopiperazine-1-carboxylate.

However, none of the tested compounds demonstrated significant anti-biofilm activity, suggesting that the observed activity of PIPC and CPZ is not attributable solely to the 2,3-DKP structure.

### MIC and Bacterial Viability in ABS Medium

To evaluate whether ABS alters antimicrobial activity, MICs were measured in both CAMHB and ABS media. Due to the opacity of ABS, MICs could not be visually determined, prompting assessment of viable cell counts instead. A 2-log reduction or less in CFU at MIC concentrations was observed in CAMHB (Fig. 6), and similar reductions in ABS medium occurred at much lower concentrations: PIPC (8 µg/mL), CPZ (4–8 µg/mL), CAZ (1 µg/mL), MEPM (0.063 µg/mL), representing substantial decreases from MICs in CAMHB (Fig.7).

**Fig. 6.**
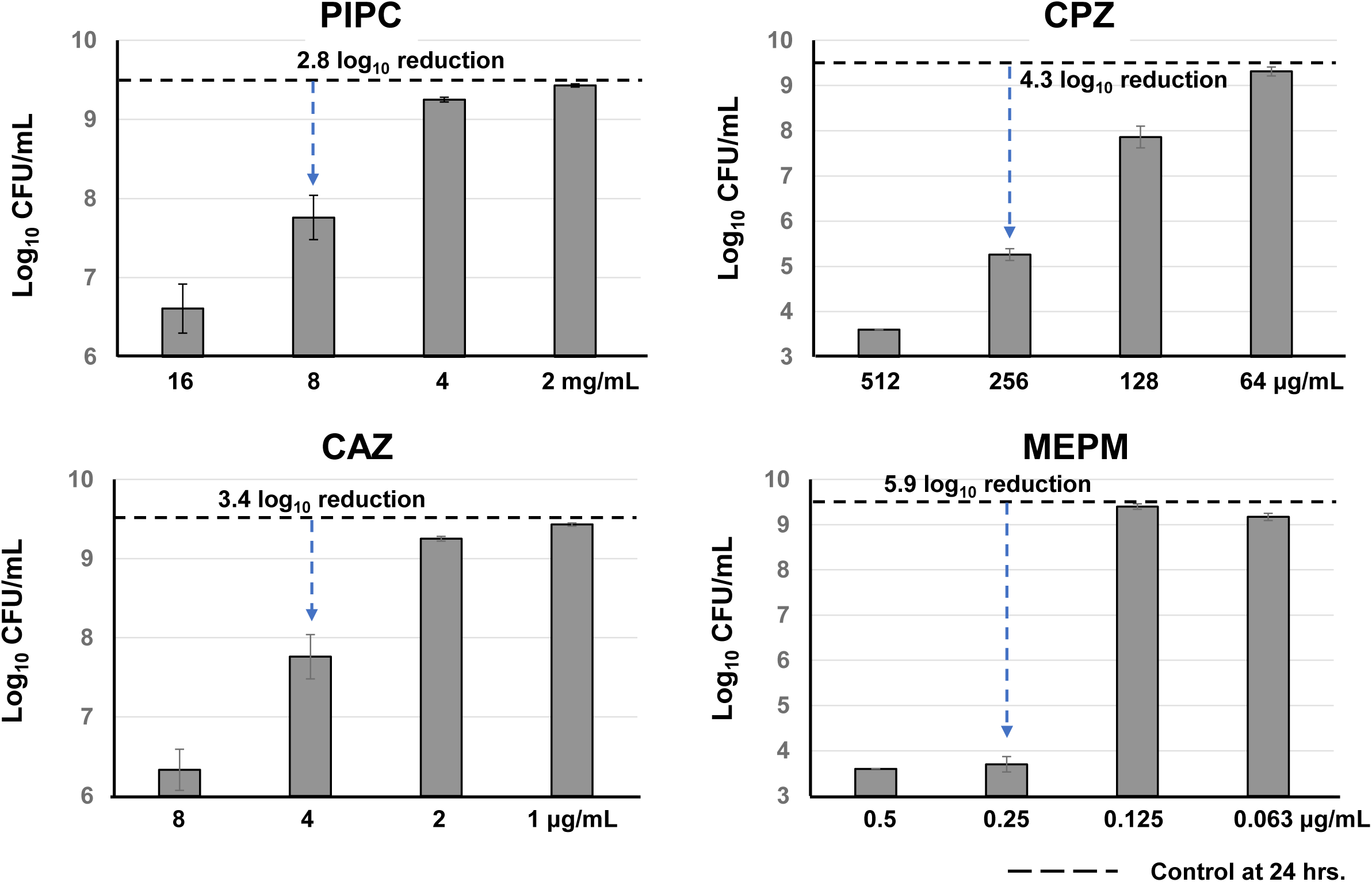
Change of CFU in MIC determination of PIPC and CAZ using CAMHB (CLSI method) in *P. aeruginosa* S3. Each experiment was evaluated in triplicate, and arrows indicate the difference between the viable counts at the MIC and the control. Each experiment was evaluated in triplicate, and expressed as average ±S.D.

**Fig. 7.**
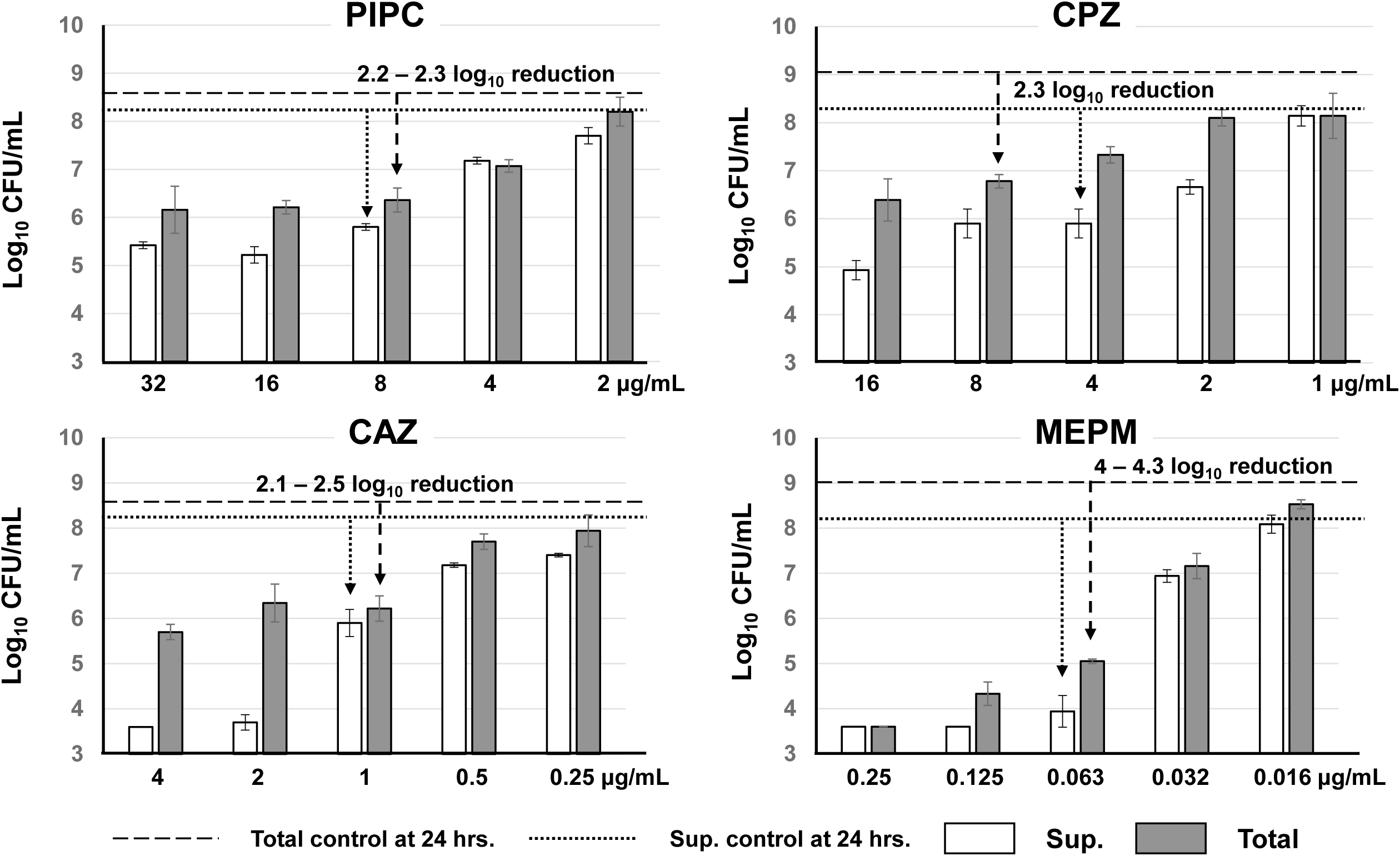
Change of colony forming unit in MIC determination of PIPC, CPZ, CAZ and MEPM using ABS medium in *P. aeruginosa* S3. Each experiment was evaluated in triplicate, and arrows indicate the difference between the viable counts at the MIC and the control at 24 hours. Each experiment was evaluated in triplicate, and expressed as average ±S.D.

Table 3 presents the MIC determined by the CLSI standard in CAMHB, the MIC measured in ABS medium, the BFIC₅₀ values in ABS, and their respective ratios. For PIPC and CPZ, MIC values decreased by 32- to 2048-fold in ABS compared to CAMHB, whereas CAZ and MEPM exhibited only a fourfold reduction. Notably, the BFIC₅₀ values were approximately one-quarter of the MICs in ABS across all four agents, indicating that inhibition of biofilm formation in ABS is likely attributable to general growth suppression rather than specific anti-biofilm mechanisms at sub-inhibitory concentrations.

**Table 3.** MIC and BFIC₅₀ values in CAMHB and ABS. The B/C (MIC/BFIC₅₀ in ABS) ratios ranged between 4 and 8, indicating that biofilm inhibition occurred at sub-MIC levels across all agents tested.

| Agents | MIC (µg/mL) |  | BFIC<sub>50</sub>*<br>(µg/mL)<br>(C) | A / B<br>ratio | A / C<br>ratio | B / C<br>ratio |
| --- | --- | --- | --- | --- | --- | --- |
|  | CAMHB<br>(A) | ABS<br>medium (B) |  |  |  |  |
| PIPC | 16384 | 8 | 2 | 2048 | 8192 | 4 |
| CPZ | 256 | 4 - 8 | 1 | 32 - 64 | 256 | 4 - 8 |
| CAZ | 4 | 1 | 0.25 | 4 | 16 | 4 |
| MEPM | 0.25 | 0.063 | 0.016 | 4 | 16 | 4 |

\*: Biofilm inhibitory concentration 50%

### Role of Serum Lipid Components

Since ABS contains lipids that can influence bacterial membrane stability, we defatted ABS using n-hexane and tested PIPC’s activity. The biofilm inhibitory effect of PIPC was reduced by 32-fold in defatted-ABS (Fig. 8), suggesting that lipophilic serum components significantly contribute to enhanced antibiotic efficacy.

**Fig. 8.**
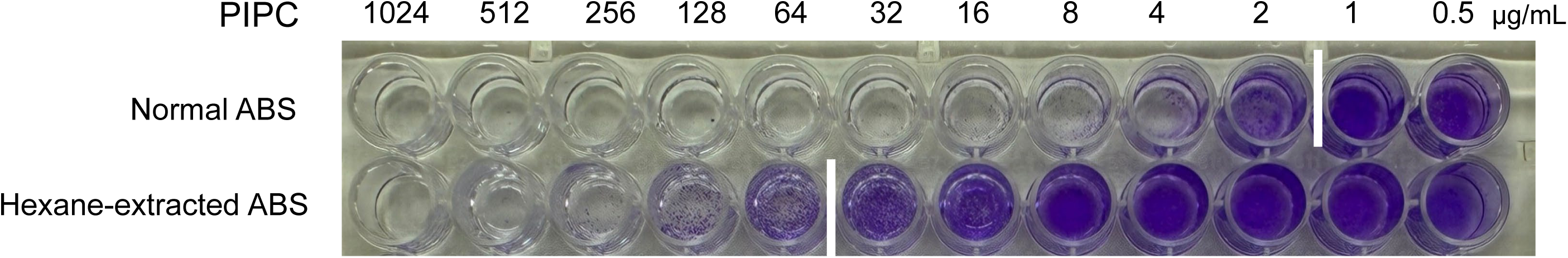
Effect of n-hexane extraction of ABS for inhibition of PIPC for biofilm formation in *P. aeruginosa* strain S3

### Effect of Divalent Cations

Supplementation of ABS medium with MgSO_4_ at concentrations of 25 mM or 100 mM significantly mitigated the growth-inhibitory effects of PIPC and CPZ, when tested at 4 times the minimum inhibitory concentration (MIC) (*p* < 0.0001). In contrast, the presence of MgSO₄ exerted minimal modulatory effects on the activity of CAZ and meropenem MEPM under identical conditions (Fig.9). These findings suggest that Mg²⁺ may counteract antibiotic-induced membrane destabilization, potentially by enhancing outer membrane stability through stabilization of lipopolysaccharide (LPS) structures. Calcium was not included in the analysis due to its propensity to form insoluble precipitates with phosphate present in the medium.

**Fig. 9.**
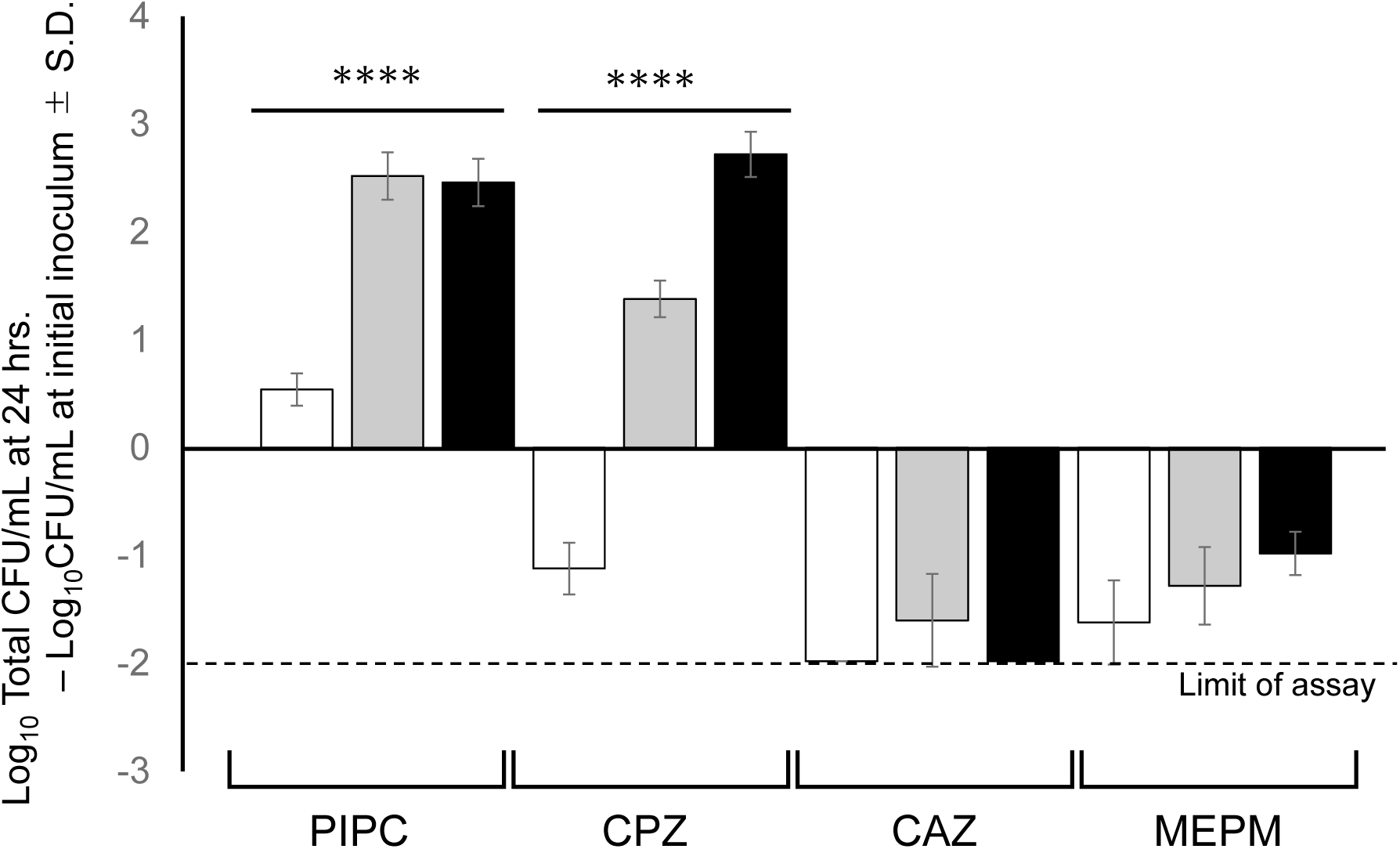
Effect of magnesium ion on antibacterial activity of PIPC, CPZ, CAZ and MEPM in ABS medium. Each drug concentration was 4 times the MIC in ABS medium, with 25 mM (grey bar) and 100 mM (black bar) magnesium ions added. After 24 hours of incubation, total cell numbers were determined by homogenization of each well with a sterilized small cotton swab and the viable bacterial count was determined. Each experiment was evaluated in triplicate, and expressed as average ±S.D. ****: *p*<0.0001

### Change of VCM susceptibility in ABS medium

Since the antimicrobial activities of PIPC and CPZ were markedly influenced by the addition of magnesium ions, it was hypothesized that lipophilic components in ABS medium chelate divalent cations, thereby destabilizing the outer membrane integrity of Gram-negative bacteria. To further investigate this membrane-disruptive effect, we evaluated the MIC of vancomycin in ABS medium. Although vancomycin typically exhibits poor activity against *P. aeruginosa* due to its inability to permeate the outer membrane, it has been shown to gain antimicrobial activity under conditions such as lipopolysaccharide (LPS) deficiency [13].

Initial MIC determination and time-kill assays conducted in CAMHB demonstrated that vancomycin exhibited a MIC of 8192 μg/mL (8 mg/mL) and reduced viable bacterial counts by approximately 4 log₁₀ after 24 hours of exposure. In contrast, when tested in ABS medium, vancomycin induced a 2 to 3 log_10_ reduction in viable cell counts at concentrations of 128 to 256 μg/mL, corresponding to a 32- to 64-fold decrease in MIC compared to CAMHB. These findings suggest that divalent cation chelation in ABS medium compromises the integrity of the bacterial outer membrane—likely by destabilizing lipopolysaccharide (LPS) structure—thereby facilitating vancomycin penetration and enhancing its bactericidal activity (Fig. 10).

**Fig. 10.**
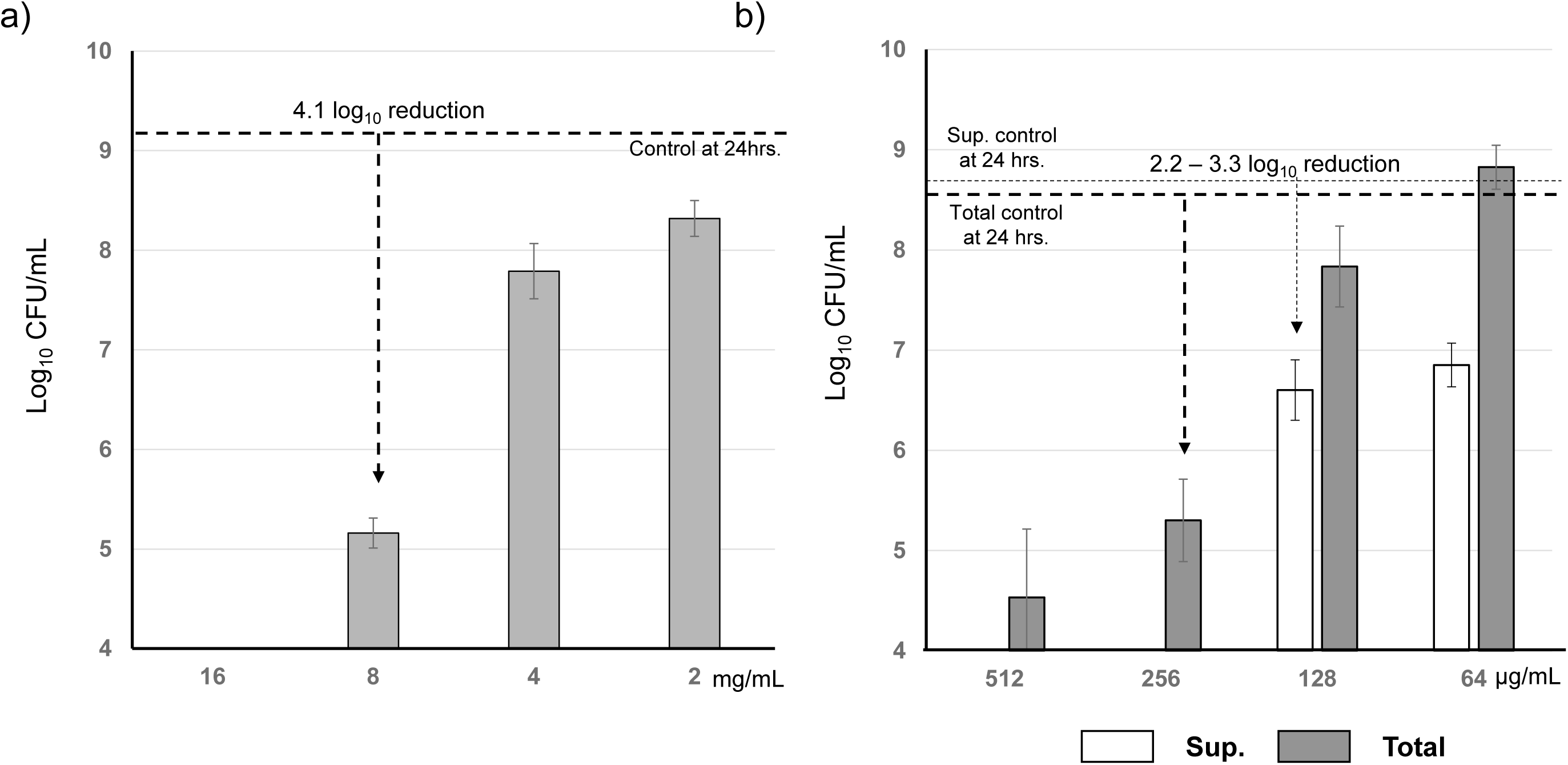
Change of CFU in MIC determination of VCM using CAMHB (a)) and ABS medium (b) in *P. aeruginosa* S3. Each experiment was evaluated in triplicate, and arrows indicate the difference between the viable counts at the MIC and the control at 24 hours. Each experiment was evaluated in triplicate, and expressed as average ±S.D.

## Discussion

This study demonstrated that certain clinical isolates of *P. aeruginosa*, particularly strain S3, rapidly and robustly form mature biofilms in ABS-containing media. Because ABS include host-derived components such as serum proteins and lipids, it may more closely mimic aspects of the *in vivo* environment than conventional laboratory media, thereby providing a potentially more relevant platform for evaluating antimicrobial efficacy under host-like conditions.

Among the β-lactam antibiotics tested, PIPC and CPZ exhibited notable anti-biofilm activity at sub-MIC concentrations in ABS medium. These antibiotics share a 2,3-diketopiperazine (2,3-DKP) core structure; however, structurally related 2,3-DKP analogs tested in this study did not exhibit comparable activity. While this observation suggests that the 2,3-DKP scaffold alone is unlikely to be sufficient for biofilm inhibition, the limited number of analogs examined—only six compounds in total—precludes a definitive conclusion regarding the role of this moiety.

Notably, previous reports have shown that 2,5-diketopiperazine modulates the LuxR-type quorum sensing (QS) system in various bacterial species and may inhibit biofilm formation by blocking this modulator function [14]. Based on structural similarity, we hypothesized that 2,3-DKPs might exert similar QS-inhibitory effects. However, under the specific conditions tested in this study, our results do not support a major role for the conserved 2,3-DKP structure in QS modulation. Nonetheless, given the limited scope of compounds and assay conditions used, we cannot exclude the possibility that certain 2,3-DKP derivatives might influence QS pathways under different environmental or genetic contexts. Additional molecular features, such as specific side chains or electronic properties, may be critical for activity. Further structure–activity relationship (SAR) studies involving a broader array of 2,3-DKP derivatives are therefore necessary.

Our findings also suggest that n-hexane–extractable serum lipids may play a role in enhancing the antimicrobial activity of certain antibiotics. These lipids are hypothesized to chelate divalent cations—particularly Mg²⁺—associated with the bacterial outer membrane, potentially leading to destabilization of the membrane structure and increased permeability to antibiotics such as PIPC and CPZ. This model is supported by the observed reduction in antimicrobial activity in defatted serum and its partial restoration upon Mg²⁺ supplementation. However, alternative or additional mechanisms—such as interactions with outer membrane proteins, modifications of lipid A, or changes in efflux pump function—may also contribute and should be examined in future work.

The reduction in the MIC of vancomycin (VCM) observed in ABS medium may serve as indirect evidence of increased outer membrane permeability. Yet, since VCM primarily inhibits peptidoglycan synthesis at the level of the inner cell wall, it is possible that other factors, such as nutrient limitations or stress responses induced by ABS components, also influence its activity. To better understand the mechanistic basis of this phenomenon, future experiments should incorporate temporal analyses of lipopolysaccharide (LPS) release and other indicators of membrane integrity and cellular stress.

These observations may align with findings by Krogfelt *et al*.[15], who reported that the phospholipid monopalmitoylphosphatidic acid (MPPA) enhances the efficacy of β-lactam antibiotics by chelating membrane-associated cations. Although the specific lipid species in ABS have not been identified, the similar enhancement of antibiotic activity seen in our study suggests that comparable mechanisms such as cation sequestration and membrane disruption may be at play.

Supporting this model, a study by Yoshimura and Nikaido [16] using proteoliposomes reconstituted with *Escherichia coli* OmpF and OmpC porins demonstrated that CPZ exhibited significantly lower porin permeability than predicted by its hydrophilicity. Penicillin-class antibiotics, including PIPC and ampicillin, are believed to cross the outer membrane predominantly via non-porin-mediated pathways [17,18]. Given the relatively large molecular size and hydrophobic side chains PIPC and CPZ, it is likely that they primarily cross the outer membrane through non-porin-mediated mechanisms. The presence of lipid-altered conditions in ABS may further facilitate such alternative diffusion pathways (Fig.11).

**Fig. 11.**
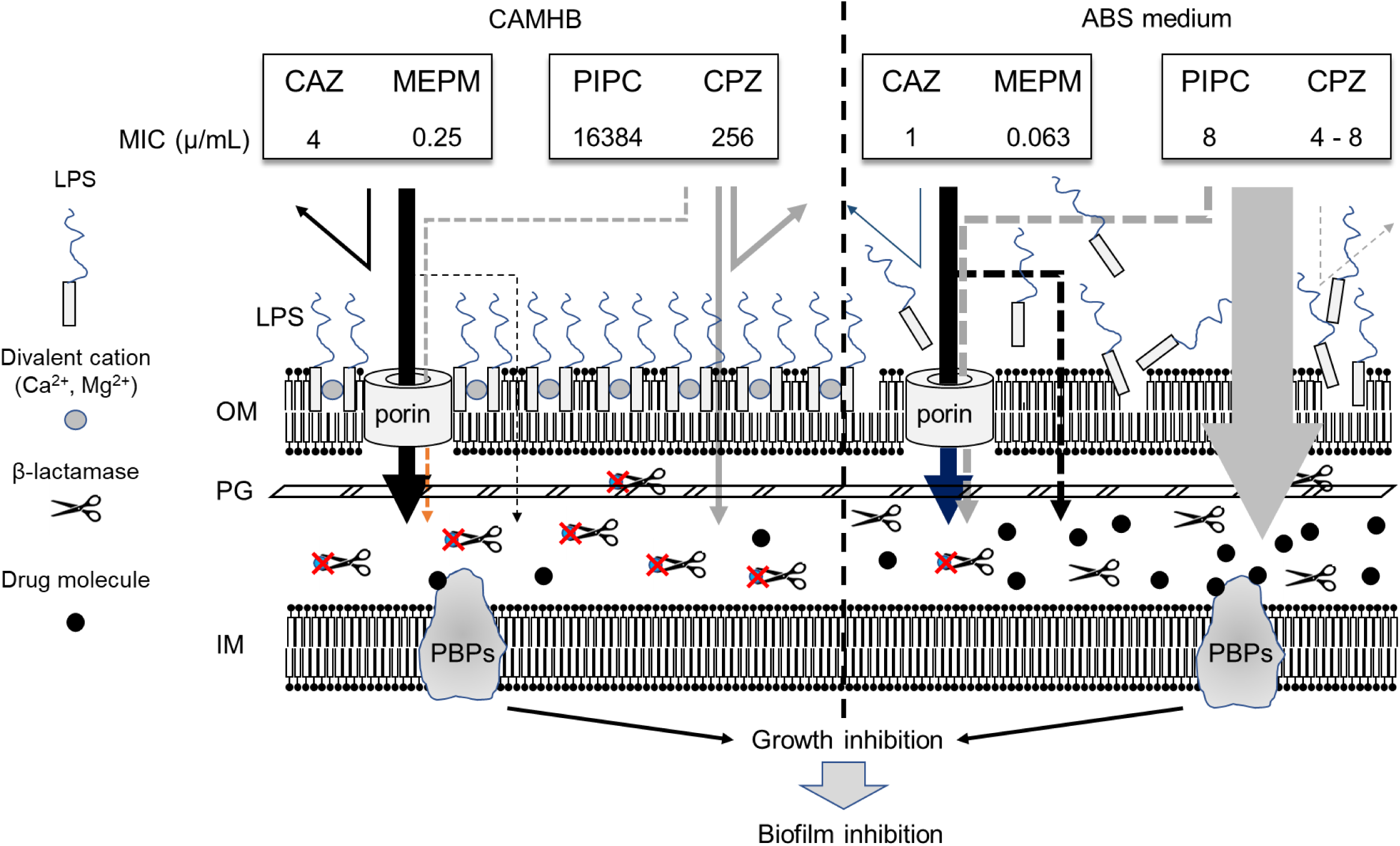
Schema of the difference in MIC and outer membrane permeability in CAMHB and ABS media. The n-hexane soluble component in ABS chelates the divalent cations of LPS and destabilizes the outer membrane structure, allowing PIPC and CPZ to permeate the outer membrane via the phospholipid bilayer. LPS: lipopolysaccharide, OM: outer membrane, PG: peptidoglycan, IM: inner membrane, PBPs: penicillin-binding proteins

Although PIPC and CPZ were effective in inhibiting biofilm formation at sub-MIC concentrations, the similarity in BFIC₅₀/MIC ratios among different antibiotics suggests that this effect may reflect general suppression of bacterial growth rather than specific interference with quorum-sensing pathways. Vieira *et al*. [19] previously reported that CPZ inhibits the *P. aeruginosa* pqs signaling pathway, reducing production of the QS autoinducers HHQ and PQS. However, under our experimental conditions, such QS-inhibitory effects were not clearly detected.

This discrepancy may result from strain-specific genetic variability, such as mutations in lasR or the *pqs* operon [20], or differences in culture medium composition, assay sensitivity, or incubation time. Further investigations incorporating multiple clinical isolates and standardized QS assays will be essential to clarify the extent and context-dependence of CPZ’s QS-modulating activity.

Interestingly, the laboratory strain PAO1 failed to form biofilms in serum-based media. The underlying reasons for this phenotype remain unclear and warrant further investigation. Factors such as the genetic background, LPS structure, exopolysaccharide production, QS pathways, and stress-response mechanisms should be systematically examined to determine whether PAO1 harbors strain-specific deficiencies in surface attachment, membrane integrity, or intracellular signaling. Although PAO1 is widely used as a reference strain for *P. aeruginosa*, it may not accurately reflect the behavior of clinical isolates under host-like conditions. Previous studies have documented considerable genetic and phenotypic variability among PAO1 sublines, including differences in motility, virulence, and biofilm formation [21]. In contrast, clinical isolates often possess unique genomic islands and enhanced stress-adaptive traits, leading to distinct biofilm phenotypes. Thus, exclusive reliance on PAO1 may yield misleading conclusions. To improve translational relevance, studies of antimicrobial susceptibility, biofilm formation, and pathogenicity under physiologically realistic conditions should incorporate a broader panel of representative clinical isolates—particularly those with robust biofilm-forming capacity, referred to as “hot” strains.

In conclusion, ABS medium offers a promising platform for studying antimicrobial and anti-biofilm effects under conditions that more closely approximate the *in vivo* environment. The enhanced efficacy of PIPC and CPZ observed in this study likely reflects a combination of host-derived factors, including but not limited to lipid-mediated outer membrane destabilization. Nevertheless, due to the compositional variability of serum and the limited scope of compounds and strains tested, cautious interpretation of these findings is warranted. Expanding future research to include a broader range of 2,3-DKP derivatives, clinical isolates, and physiologically relevant conditions will be critical for elucidating the complex interactions between antibiotics, bacterial physiology, and host factors.

## TRANSPARENCY DECLARATION

### Data availability

The authors confirm that the data supporting the findings of this study are available in the article.

### Declaration of Generative AI and AI-assisted technologies in the writing process

During the preparation of this work, the authors used Claude (Anthropic) in order to assist with sentence structuring and rephrasing for improved flow. The authors reviewed and edited the content as needed and take full responsibility for the content of the publication.

## References

1. Miller WR, Arias CA. 2024. ESKAPE pathogens: antimicrobial resistance, epidemiology, clinical impact and therapeutics. Nat Rev Microbiol. 22:598–616.

2. Haque M, Sartelli M, McKimm J, Bakar MA. 2018. Health care-associated infections—an overview. Infect Drug Resist. 11:2321–2333.

3. Costerton JW, Stewart PS, Greenberg EP. 1999. Bacterial biofilms: a common cause of persistent infections. Science. 284:1318–1322.

4. Flemming HC. 2016. EPS—then and now. Microorganisms. 4:41.

5. Stewart PS, Franklin MJ. 2008. Physiological heterogeneity in biofilms. Nat Rev Microbiol. 6:199–210.

6. Soares A, Alexandre K, Etienne M. 2020. Tolerance and persistence of *Pseudomonas aeruginosa* in biofilms exposed to antibiotics. Front Microbiol. 11:2057.

7. Bjarnsholt T, Ciofu O, Molin S, Givskov M, Høiby N. 2013. Applying insights from biofilm biology to drug development: can a new approach be developed? Nat Rev Drug Discov. 12:791–808.

8. Walters MC 3rd, Roe F, Bugnicourt A, Franklin MJ, Stewart PS. 2003. Contributions of antibiotic penetration, oxygen limitation, and low metabolic activity to tolerance of *Pseudomonas aeruginosa* biofilms to ciprofloxacin and tobramycin. Antimicrob Agents Chemother. 47:317–323.

9. Wijesinghe G, Dilhari A, Gayani B, Kottegoda N, Samaranayake L, Weerasekera M. 2019. Influence of laboratory culture media on *in vitro* growth, adhesion, and biofilm formation of *Pseudomonas aeruginosa* and *Staphylococcus aureus*. Med Princ Pract. 28:28–35.

10. Plano LMD, Caratozzolo M, Conoci S, Guglielmino SPP, Franco D. 2024. Impact of nutrient starvation on biofilm formation in *Pseudomonas aeruginosa*: an analysis of growth, adhesion, and spatial distribution. Antibiotics (Basel). 13:987.

11. Hammond A, Dertien J, Colmer-Hamood JA, Griswold JA, Hamood AN. 2010. Serum inhibits *Pseudomonas aeruginosa* biofilm formation on plastic surfaces and intravenous catheters. J Surg Res. 159:735–746.

12. Clinical and Laboratory Standards Institute (CLSI). 2024. M07-Ed12. Methods for dilution antimicrobial susceptibility tests for bacteria that grow aerobically, 12th ed. CLSI, Wayne, PA

13. Vidaillac C, Benichou L, Duval RE. 2012. *In vitro* synergy of colistin combinations against colistin-resistant *Acinetobacter baumannii*, *Pseudomonas aeruginosa*, and *Klebsiella pneumoniae* isolates. Antimicrob Agents Chemother. 56:4856–4861.

14. Campbell J, Lin Q, Geske GD, Blackwell HE. 2009. New and unexpected insights into the modulation of LuxR-type quorum sensing by cyclic dipeptides. ACS Chem Biol. 4:1051–1059.

15. Krogfelt KA, Utley M, Krivan HC, Laux DC, Cohen PS. 2000. Specific phospholipids enhance the activity of beta-lactam antibiotics against *Pseudomonas aeruginosa*. J Antimicrob Chemother. 46:377–384.

16. Yoshimura F, Nikaido H. 1985. Diffusion of beta-lactam antibiotics through the porin channels of *Escherichia coli* K-12. Antimicrob Agents Chemother. 27:84–92.

17. Sawai T, Hiruma R, Kawana N, Kaneko M, Taniyasu F, Inami A. 1982. Outer membrane permeation of beta-lactam antibiotics in *Escherichia coli*, *Proteus mirabilis*, and *Enterobacter cloacae*. Antimicrob Agents Chemother. 22:585–592.

18. Yamaguchi A, Hiruma R, Sawai T. 1982. Phospholipid bilayer permeability of beta-lactam antibiotics. J Antibiot (Tokyo). 35:1692–1699.

19. Vieira TF, Leitão MM, Cerqueira NMFSA, Sousa SF, Borges A, Simões M. 2024. Montelukast and CPZ act as antiquorum sensing and antibiofilm agents against *Pseudomonas aeruginosa*. J Appl Microbiol. 135:lxae088.

20. O’Connor K, Zhao CY, Mei M, Diggle SP. 2022. Frequency of quorum-sensing mutations in *Pseudomonas aeruginosa* strains isolated from different environments. Microbiology (Reading). 168:001265.

21. Chandler CE, Horspool AM, Hill PJ, Wozniak DJ, Schertzer JW, Rasko DA, Ernst RK. 2019. Genomic and phenotypic diversity among ten laboratory isolates of *Pseudomonas aeruginosa* PAO1. J Bacteriol. 201:e00595–18.

